# Revealing acquired resistance mechanisms of kinase-targeted drugs using an on-the-fly, function-site interaction fingerprint approach

**DOI:** 10.1101/831800

**Authors:** Zheng Zhao, Philip E. Bourne

**Author notes:** **Corresponding author** Philip E. Bourne.

## Abstract

Although kinase-targeted drugs have achieved significant clinical success, they are frequently subject to the limitations of drug resistance, which has become a primary vulnerability to targeted drug therapy. Therefore, deciphering resistance mechanisms is an important step in designing more efficacious, anti-resistant, drugs. Here we studied two FDA-approved kinase drugs: Crizotinib and Ceritinib, which are first- and second-generation anaplastic lymphoma kinase (ALK) targeted inhibitors, to unravel drug-resistance mechanisms. We used an on-the-fly, function-site interaction fingerprint (on-the-fly Fs-IFP) approach by combining binding free energy surface calculations with the Fs-IFPs. Establishing the potentials of mean force and monitoring the atomic-scale protein-ligand interactions, before and after the L1196M-induced drug resistance, revealed insights into drug-resistance/anti-resistant mechanisms. Crizotinib prefers to bind the wild type ALK kinase domain, whereas Ceritinib binds more favorably to the mutated ALK kinase domain, in agreement with experimental results. We determined that ALK kinase-drug interactions in the region of the front pocket are associated with drug resistance. Additionally, we find that the L1196M mutation does not simply alter the binding modes of inhibitors, but also affects the flexibility of the entire ALK kinase domain. Our work provides an understanding of the mechanisms of ALK drug resistance, confirms the usefulness of the on-the-fly Fs-IFP approach and provides a practical paradigm to study drug-resistance mechanisms in prospective drug discovery.

## Introduction

Targeted drug therapy has become a major cancer treatment beyond surgery, radiation therapy, chemotherapy and immunotherapy^1^. Validating primary targets and developing targeted drugs has received increasing attention^2^. In concert, over the last 20 years, the kinase family has been recognized as important drug targets^3–4^. As of August 2018, 42 kinase-targeted drugs have been approved by the U.S. Food and Drug Administration (FDA)^5^ and have revolutionized clinical therapy against multiple diseases, especially different cancer types, such as non-small cell lung cancer (NSCLC), melanoma and leukemia ^1, 6^. However, the efficacy of drugs is frequently subject to the limitation of drug-acquired resistance. For example, Crizotinib has been frequently found to be ineffective for the majority of patients after one to two years’ treatment against ALK-positive NSCLC, due to the acquired mutations at the binding site ^7^. As such, patients have a need for next-generation drugs effective against acquired drug resistance^5, 8^. Thus, deciphering the mechanisms of acquired resistance is a step forward in designing novel and efficacious anti-resistant drugs^9^.

Exploring resistance mechanisms has been challenging due to the diversity of specific drug-binding mechanisms. For example, Yun et al.^10^ concluded that the resistance against Gefitinib and Erlotinib, which are two FDA-approved kinase drugs used to treat patients with EGFR-overexpression-induced NSCLC, is caused by the gatekeeper T790M mutation; this mutation increased the binding affinity of ATP in EGFR kinase. In another example, the resistance against Dasatinib, a tyrosine-kinase inhibitor employed to treat people with Ph+ chronic myeloid leukemia and acute lymphoblastic leukemia^11^, is caused by weakening the drug’s affinity to the binding pocket due to a T338M mutation in the cSrc and Abl kinase family^12^. Worse still, the resistance mechanisms remain undetermined for most targeted drugs^6^. For instance, Crizotinib is the first FDA-approved drug against ALK-positive NSCLC, and its drug resistance frequently occurs in patients, along with the development of mutations to the binding site^13–14^. On one hand, researchers have suggested that the mechanism involves acquired mutations blocking the binding of inhibitors, such as where the L1196M mutation confers ALK kinase high-level resistance to Crizotinib^15^ through decreased binding affinity^16^. In contradiction, the high-resolution crystal structures of the wild-type and L1196M ALK-Crizotinib complexes (PDB ids: 2xp2 and 2yfx) suggest the binding modes are similar^17^. Thus, it is essential to develop efficient strategies to reveal resistance mechanisms when designing future drugs. To this end, we here provide an effective method called the on-the-fly, function-site interaction fingerprint (Fs-IFP) approach to explore kinase drug-resistance mechanisms.

The on-the-fly Fs-IFP approach we propose is formed by combining binding free-energy calculations with the Fs-IFP encoding. The binding free-energy calculation is an important measure in understanding the ligand-binding recognition processes and differs from the binding affinity for ligands^18^. Theoretically, there are different methods to calculate binding free energy, such as thermodynamic integration, free-energy perturbation and probability distributions and histograms^19^. Here, we used a sophisticated, improved sampling strategy (umbrella sampling) with a WHAM analysis to obtain the binding free-energy surface (FES), which has been successfully applied in different complex systems^20^. More importantly, in our proposed scheme, we monitored the binding process by using an Fs-IFP encoding strategy. This strategy has practical applications in analyzing a variety of binding pockets^21–24^ to reveal details of binding, notably the differences before and after mutation(s).

**Figure 1.**
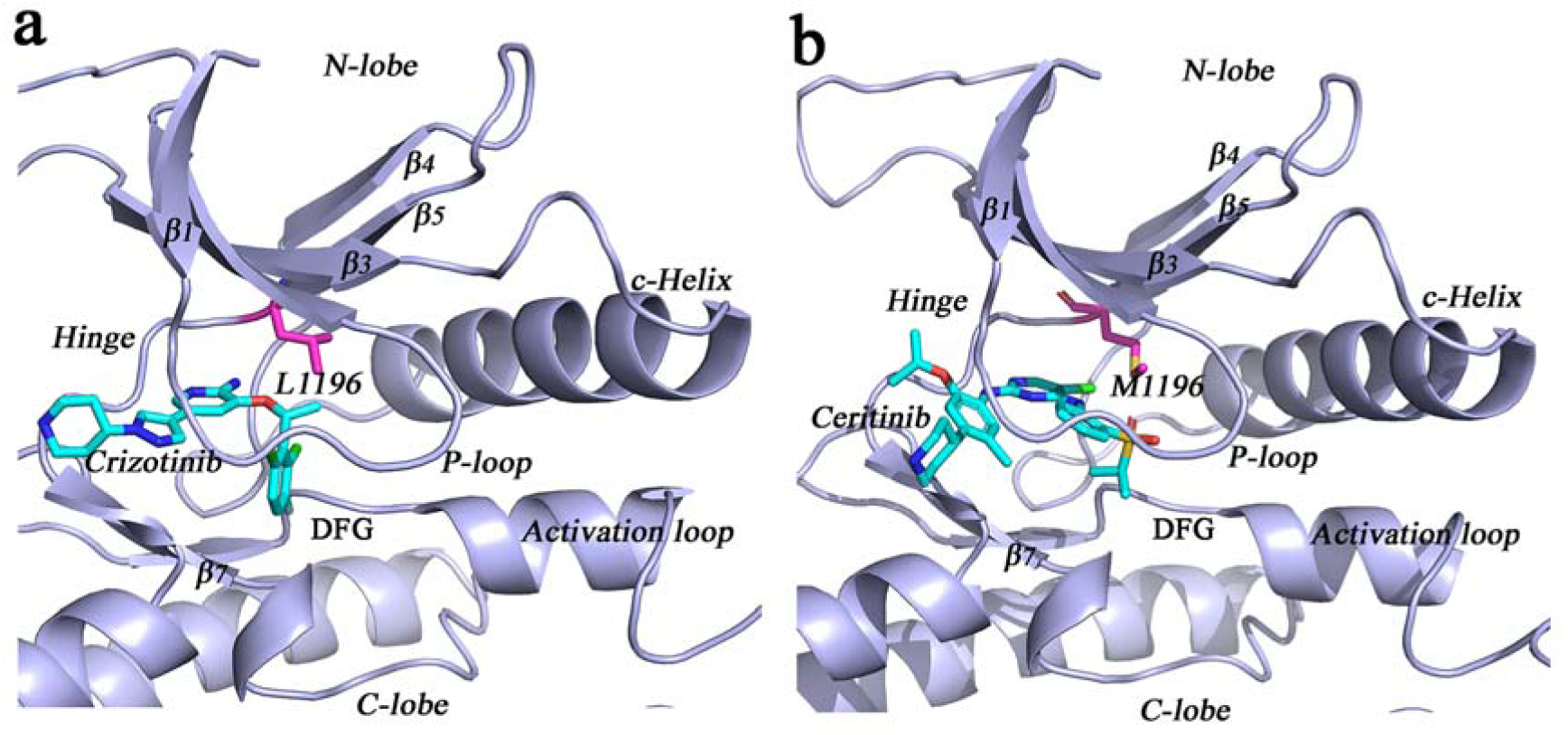
Representative ALK-drug interactions. (a) Crizotinib-bound ALK complex. (b) Ceritinib-bound ALK complex. Residue 1196 is the gatekeeper, and the L1196M mutation is shown when binding Ceritinib.

In this work, we explore the resistance/anti-resistant mechanisms of two drugs (Crizotinib and Ceritinib) against ALK in the case of wild type and L1196M mutation. Ceritinib is a second-generation ALK inhibitor which efficiently overcomes L1196M-induced drug resistance. Ceritinib has shown stronger inhibition, not only for the wild-type ALK, but also for the L1196M mutant^25^. Using the on-the-fly Fs-IFP approach, we obtained the dynamic binding details of the two drugs by exploring the binding free energy surfaces (FESs). We revealed the details of a resistance mechanism for Crizotinib as well the anti-resistant mechanism of Ceritinib. This includes details of the flexibility of the apo and holo ALK kinase domains. The hope is that these insights contribute to an understanding of the drug resistance mechanisms and to next-generation drug design. The on-the-fly pipeline for studying drug resistance mechanisms can be applied more generally beyond ALK kinase.

## Results

### Drug binding profiles

We determined the binding processes of Crizotinib and Ceritinib by exploring the corresponding binding free-energy surfaces (Figure 2). We note that every binding free-energy profile for different drug-binding systems is an energy-decreasing procedure from the dissociated state to the bound state, along the predefined reaction coordinates (Supplemental Figure S1); that means the drug-binding process is spontaneous and thermodynamically favorable. In the bound state, the potentials of mean force (PMF) have a global minimum state, and the relative PMF well depths are −9.8±0.7, −11.0±0.5, −11.3±0.7, and −13.6±0.8 kcal.mol^−1^ for wild/mutated Crizotinib-bound and Ceritinib-bound systems, respectively, in agreement with experimental results^16, 26^. Based on the comparisons of wild/mutated PMFs (Figure 2a), Crizotinib prefers to bind to the wild ALK kinase binding site rather than the L1196M-mutated binding site. Comparatively, Ceritinib binds more favorable to a L1196M-mutated ALK kinase (Figure 2b). Moreover, Ceritinib has slightly lower binding free-energy potential, which means that Ceritinib binds not only to wild-type but also to mutated ALK binding sites more closely than Crizotinib. Given this information, we explore the resistance mechanisms of Crizotinib, and the anti-resistant mechanisms of Ceritinib at the atomic level using the Fs-IFP encoding method.

**Figure 2.**
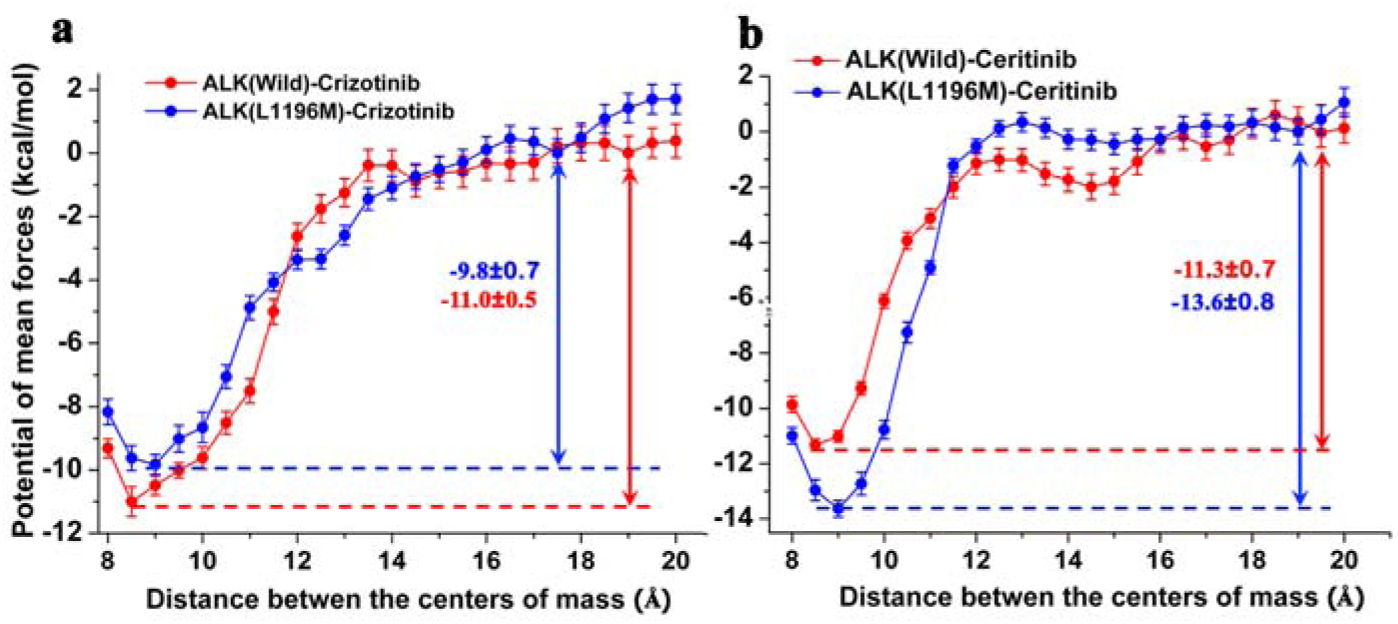
Potential of mean force profiles for wild-type or mutated drug-binding processes along with the predefined reaction coordinates. (a) Crizotinib binding profile; (b) Ceritinib binding profile.

### Drug-binding features

The Fs-IFPs for the structures of the drug-bound complexes illustrate the binding characteristics and differences before and after mutation. We extracted Fs-IFPs for conformations of the sampling windows with the lowest PMF (i.e., the bound state). Significant differences are presented (Figure 3).

**Figure 3.**
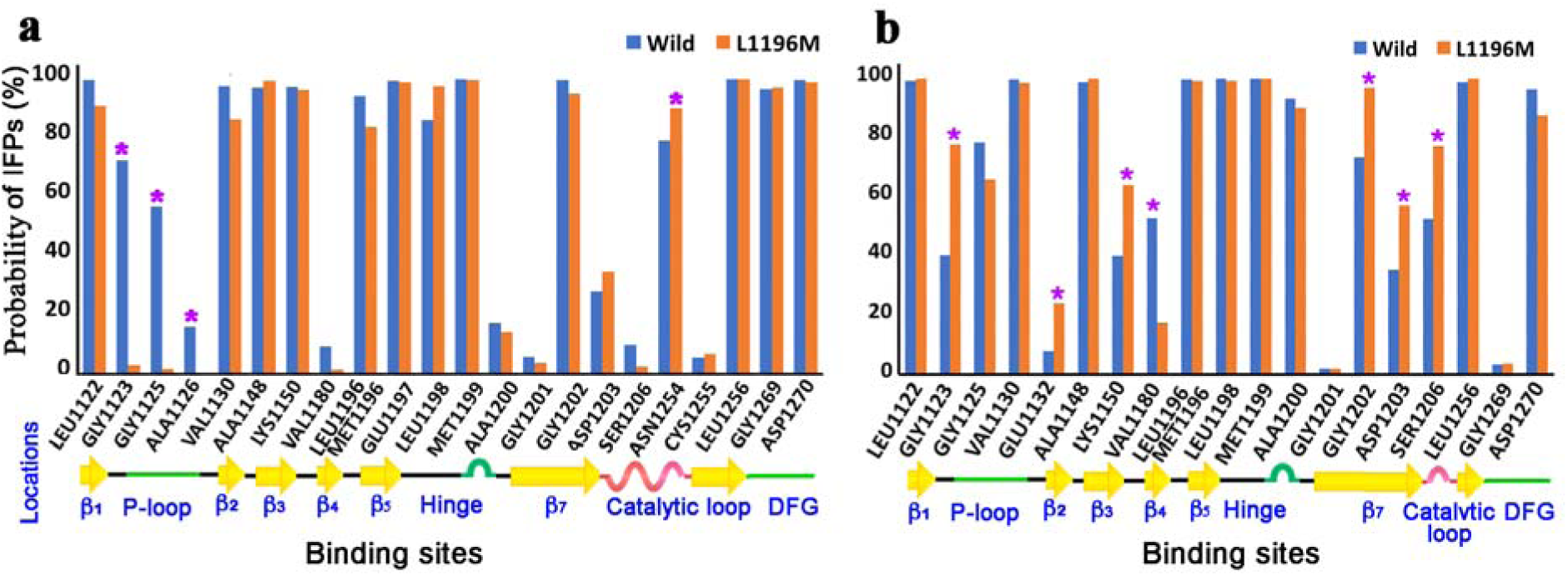
Comparisons of the Fs-IFPs between the wild-type and the L1196M mutated systems in the drug-bound states. (a) Crizotinib-bound systems. (b) Ceritinib-bound systems.

In total, the features of Crizotinib binding to the wild or mutated ALK kinase are very similar (Figure 3a). At the regions of β_1_, β_2_, β_3_, β_5_, Hinge, β_7_ and DFG, there are conserved ALK-Crizotinib interactions (these interactions occur in more than 90% of conformations), which show no significant changes before or after mutation. Noteworthy is the lack of significant change in the hydrophobic interactions between residue 1196 and Crizotinib regardless of the mutation. Further, at the P-loop region consisting of G1123, G1125 and A1126, the interactions are significantly weakened in the mutated systems (Figure 3a). These decreased interactions will induce the P-loop to become more flexible and more sensitive to the selectivity of Crizotinib, thereby inferring drug resistance.

Similarly, the conserved ALK-Ceritinib interactions also exist in the regions of β_1_, β_3_, β_5_, Hinge, Catalytic loop, and DFG, regardless of the L1196M mutation (Figure 3b). However, in the mutated case, there are multiple residues including G1123, E1132, K1150, G1202, D1203, and S1206, presenting more conserved interactions than in wild type. These more conserved interactions contribute to the binding affinity of Ceritinib in the mutated system as shown in the corresponding free-energy profile (Figure 2b). It is worth noting that G1202, D1203, and S1206 are located in the region of β7, which constitutes the front pocket^21^. Meanwhile, G1123 is located in the region of the P-loop; the P-loop also constitutes part of the front pocket. Thus, these more conserved interactions at the front pocket contribute to overcome drug resistance. Compared to the change in ALK-Crizotinib interaction characteristics before and after mutation, the front pocket, including P-loop and β7, plays the key role in inducing or overcoming the acquired L1196M drug resistance. This suggests an anti-resistant drug should be designed with more attention to achieving conserved interactions in the region of the front pocket.

### Hydrophobic interactions at the gatekeeper

The mutation of the gatekeeper (residue 1196) is often attributed to drug resistance^12^. Here we carefully checked the hydrophobic interactions between the gatekeeper and the ligand before and after mutation in the bound state. We present the distribution of the hydrophobic interactions between the gatekeeper and the ligand (Figure 4). For the wild-type ALK-Crizotinib system, the maximum probability distribution (MPD) is 3.9 Å, where there are stronger hydrophobic interactions compared to the L1196M-mutated system, which has an MPD of 4.2 Å (Figure 4a). In contrast, in the Ceritinib-bound case, there is a stronger hydrophobic interaction in the L1196M system, with an MPD of 3.55 Å, than in the wild-type with an MPD of 3.7 Å. These trends in hydrophobic interaction are in agreement with the drug-binding affinities implying that the hydrophobic interactions at the gatekeeper also play a role in ligand binding. Thus, along with the Fs-IFP features, the hydrophobic interactions result in slight drug resistance to Crizotinib and contribute to overcome the drug resistance to Ceritinib.

**Figure 4.**
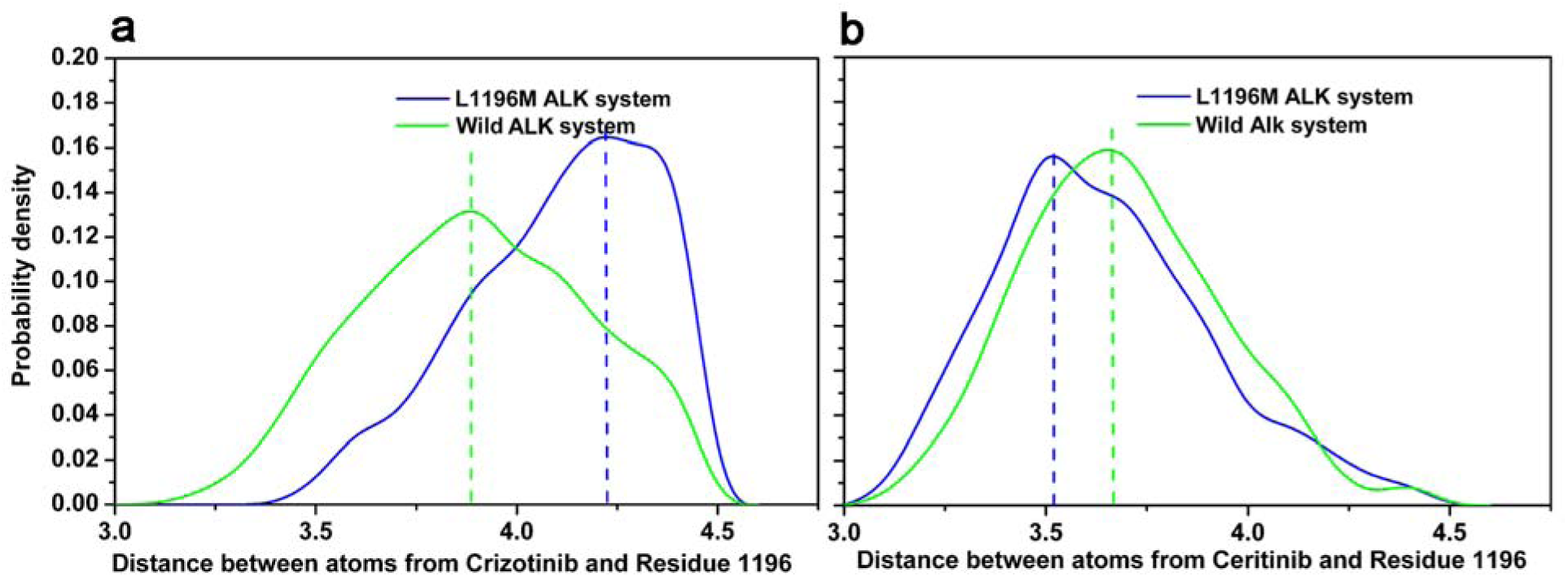
Probability distribution of interatomic distances between Residue 1196 and the corresponding ligand. (a) in the Crizotinib-bound system. (b) in the Ceritinib-bound system.

### Collective motions of the ALK kinase domain

The collective change in conformation of target proteins always leads to drug-resistance mechanisms^27–28^. We analyzed the trajectories of both wild-type and mutant apo ALK kinase conformations using principal component analysis (PCA) (Figure 5a). Overall the dynamics trajectories show similar features and magnitudes in the core structures of both wild-type and mutated apo AKL kinase domains. However, in regions of flexibility, the collective motions of the apo AKL kinase domain indicate a significant difference before and after the L1196M mutation (Figure 5a). The catalytic loop presents large flexibility, with a strong dynamic magnitude in the wild type, whereas the N-lobe has increased flexibility in the mutated kinase domain (Figure 5a). The difference in mutation-induced flexibility would, therefore, seem a possible contributor to drug resistance.

**Figure 5.**
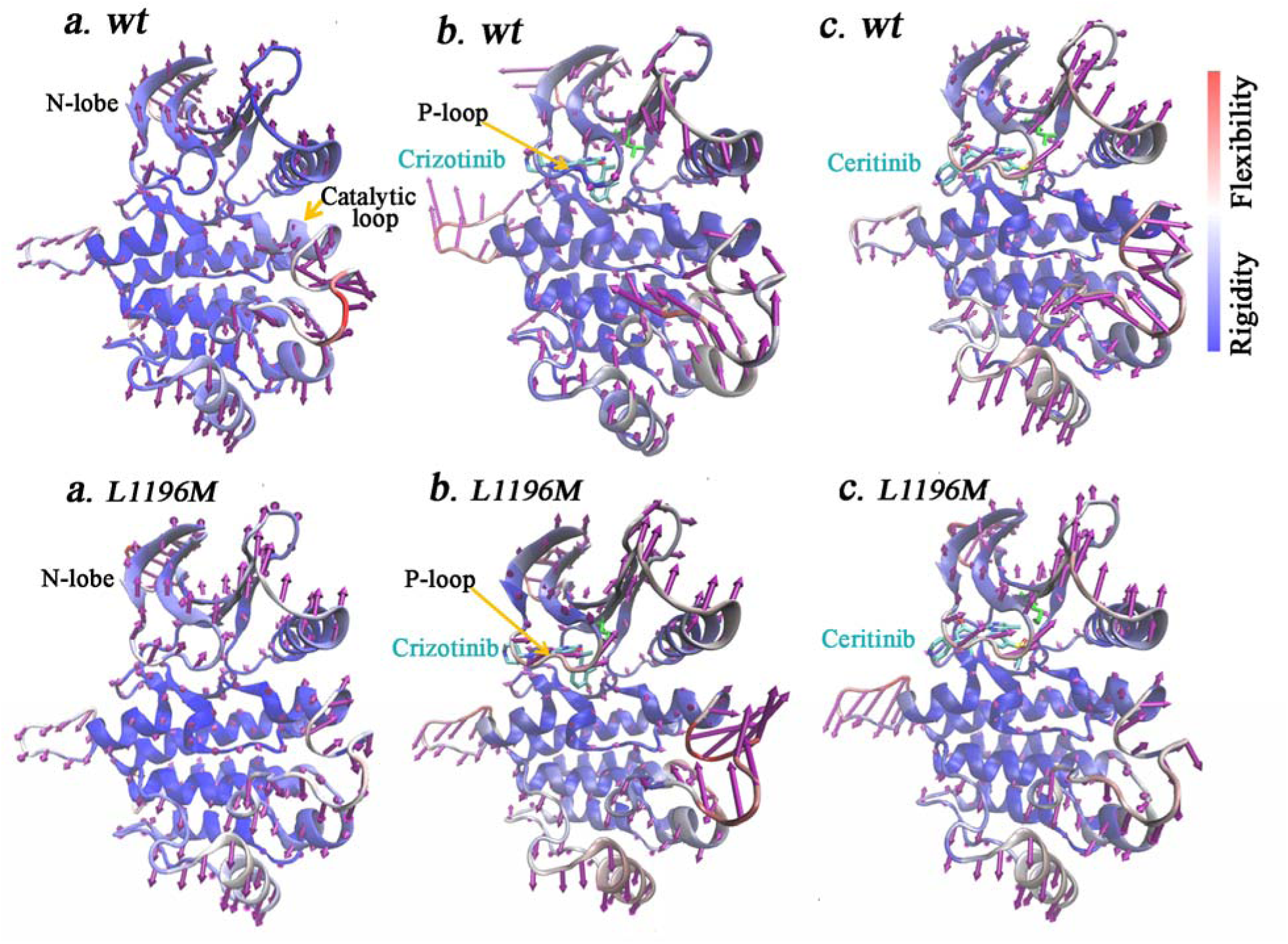
Essential dynamics analysis using PCA. (a) on the apo ALK kinase domain system; (b) on the Crizotinib-bound ALK system; (c) on the Ceritinib-bound ALK system. Arrows (purple) on every structure show the first principal direction for each residue with significant perturbation. The length of each arrow shows the magnitude of the motion for each residue. Different backbone colors show structural flexibility (red) or rigidity (blue).

Considering the ALK-drug complexes with the lowest PMF, in the wild-type ALK-Crizotinib complex (Figure 5b), Crizotinib is located at the binding pocket, competes with ATP and has a stable binding interaction with ALK. Compared to apo ALK, the essential dynamics of wild-type Crizotinib-bound ALK kinase shows a similar magnitude but moves in a different direction. However, in the mutated ALK-Crizotinib system, the conformation space appears to be similar to that in the L1196M apo ALK system (Figure 5a-b, L1196M). Clearly, the essential motion is different between the wild-type and mutated ALK-Crizotinib complexes, notably the flexibility of the P-loop. In the wild type, the P-loop is rigid, and interacts with Crizotinib, but appears to be more flexible in the mutated case (Figure 5b). That behavior agrees with the observed differences in the interaction fingerprints (Figure 3).

To evaluate the population propensity of Ceritinib-bound ALK systems, we also analyzed the conformations from the trajectories of the ALK-Ceritinib complexes (Figure 5c). Here, conformational motions are distinctly different from the Crizotinib-bound wild-type system, but similar to the apo kinase, regardless of whether they are wild-type or mutated. There is a slight propensity for flexibility at the P-loop region in L1196M Ceritinib-bound complexes, which is in accordance with the dynamic modes of the apo ALK kinase, which is also more flexible at the N-lobe, due to mutations.

In summary, the PCA analysis supports the difference in Fs-IFP’s before and after mutations, and demonstrates the dynamic motions of the apo and holo kinase domains. The Crizotinib-bound kinase domain has a different dynamic mode from the apo ALK conformations or the Ceritinib-bound ALK complexes.

## Conclusions

We present an on-the-fly Fs-IFP approach by combining free-energy calculations and Fs-IFP encoding to reveal the resistance mechanism for two kinase-targeted drugs. Our calculations support experimental evidence that Crizotinib induces drug resistance, whereas Ceritinib can overcome L1196M-induced drug resistance. Fs-IFP encoding reveals the details of the binding mode before and after mutation, which is useful in differentiating subtle changes in the target-drug interaction. This finding has broader implications across the kinase family given the high similarity of ATP binding. We also studied the dynamic modes for the ALK kinase domain and the drug-bound kinase complexes. The wild-type Crizotinib-bound ALK complex displays a different motion than the wild-type or mutated apo ALK kinase domain. In comparison, the Ceritinib-bound complex has a dynamic mode similar to apo ALK kinase, which is favorable for overcoming drug resistance.

Our computational analysis provides detailed insights into experimentally observed first-generation drug resistance and second-generation drug anti-resistance. It is challenging to highlight the key factors responsible for ALK-targeted drug resistance because there are conserved interactions between ALK kinase and the drug (both Crizotinib and Ceritinib), especially, in the regions of the Hinge and the gatekeeper. Using an on-the fly Fs-IFP scheme, we not only present the binding characteristics of every ligand at the binding sites but also exhibit the specific change in Fs-IFPs after mutation. The difference in Fs-IFPs shows that the ALK kinase-drug interactions in the region of the front pocket are related to drug resistance. Overall this methodology paves the way for designing next-generation anti-resistant drugs.

## Methods

### 1. The on-the-fly function-site interaction fingerprint (on-the-fly Fs-IFP) approach

In this work, we provided an on-the-fly Fs-IFP approach by combining the binding free energy surface calculation and the function-site interaction fingerprint method (Fs-IFP).

#### 1.1 Calculating the binding free energy surface

The binding free energy surface is calculated by using umbrella sampling (US) with the weighted histogram analysis method (WHAM). The US method represents significant progress in enhancing the sampling of the free energy surface^29^. The theoretical basis has been well described ^29–30^ and not repeated here. Following US, using WHAM for free energy calculations has emerged as an efficient scheme^20^, especially as the number of dimension of the reaction coordinates and the complexity of free energy surface increases^31^. A brief overview of the WHAM extension is provided in the supporting information.

#### 1.2 Encoding function-site interaction fingerprints (Fs-IFPs)

Fs-IFP is a method to determine protein-ligand interaction characteristics at the functional site and to do so on a proteome-wide scale as detailed in previous applications^21–23^. The protein– ligand interfacial interaction is described using 1D fingerprints, which can discriminate between the large number of potential ligand binding modes.

The method has been described in detail elsewhere^21–23^. Briefly, we first aligned all of binding sites with a sequence-independent alignment tool SMAP using default parameters^32^. Then, for every conformation, the protein-ligand interactions of every involved residue are encoded as a 7-bit fingerprint using the predefined geometric rules^32^ for seven types of interactions: (1) van der Waals; (2) aromatic face to face; (3) aromatic edge to face; (4) hydrogen bond (protein as hydrogen bond donor); (5) hydrogen bond (protein as hydrogen bond acceptor); (6) electrostatic interaction (protein positively charged); and (7) electrostatic interaction)^33^. Output includes the name and index of the interacting atoms corresponding to the fingerprints, in other words, which atoms are contributing to specific interaction features.

In this way, we systematically obtained the interaction features by encoding the Fs-IFP of every snapshot through every umbrella sampling trajectory.

### 2. Computational details

Initial conformations of the wild-type and the L1196M mutated ALK-Crizotinib complex were taken from the Protein Data Bank^34^ (PDB IDs: 2xp2^35^ and 2yfx^17^, respectively). Moreover, the two PDB structures after removing the corresponding ligand were used as the initial structures for the wild-type and mutant apo ALK kinase systems, respectively. The missing loops were built using the Modeller software^36^ using the corresponding ALK amino acid sequence^37^. Similarly, the wildtype ALK-Ceritinib-binding complex was taken from the PDB (PDB ID: 4mkc). Because no L1196M ALK-Ceritinib-binding complex was available, we obtained the complex by docking Ceritinib into an ALK L1196M conformation (PDB ID: 2yfx) using Surflex v4.1^38^. The force field parameters and the topology files of Crizotinib and Ceritinib were assignment using the ParamChem server^39–40^. The protonation states of the charged residues of all six systems were determined assuming a constant pH of 7.0. Then all the systems were solved in a rectangular water box with an 18Å buffer from any solute atom, respectively. Counter ions were added to ensure an ion concentration of 0.20 M and electroneutrality. The CHARMM36 force field^41^, CHARMM general force field^42^ and TIP3P force field^43^ were used for the ALK, Crizotinib and Ceritinib and water molecules, respectively. All systems were optimized using an ACEMD MD setup protocol^44^: 2ps minimization, 100ps for NVT, 1ns for NPT with heavy-atom constraints and 1ns for NPT without any constraints. Subsequently, 150ns MD simulations were performed to equilibrate every system. In all MD simulations, all bonds were constrained using SHAKE and the integration time step was 4 fs. The temperature bath used the Langevin method, and 1atm pressure was maintained using the Berendsen method^45^. All MD simulations and later umbrella sampling were carried out using the ACEMD software^44^.

The free energy simulation of ALK-drug system was carried out in a serial of discrete windows. For every ALK-drug system, we used 25 windows following the bias potential with the distance of centers of mass as the predefined reaction coordinates (supplemental information). In every window, 160ns MD simulations were carried out and the last 100ns MD trajectories were analyzed to obtain PMFs and encode the Fs-IFPs.

For the apo ALK kinase, a 1200ns MD simulation was carried out and the last 1000ns trajectory used as the essential dynamics analysis. All the essential dynamics analysis was performed using PCA and visualized using Normal mode Wizard in VMD^46^.

The Fs-IFP encoding was run using IChem software^47^ with the similarity of the pairwise Fs-IFPs calculated using the Tanimoto coefficient (TC)^48^.

## Acknowledgements

We thank Prof. Ajay N. Jain for providing the Surflex v4.1. We thank Prof. Haiyan Liu for his guidance in the FES analysis using WHAM. This work was partly supported by the University of Virginia (PEB).

## Table of Content

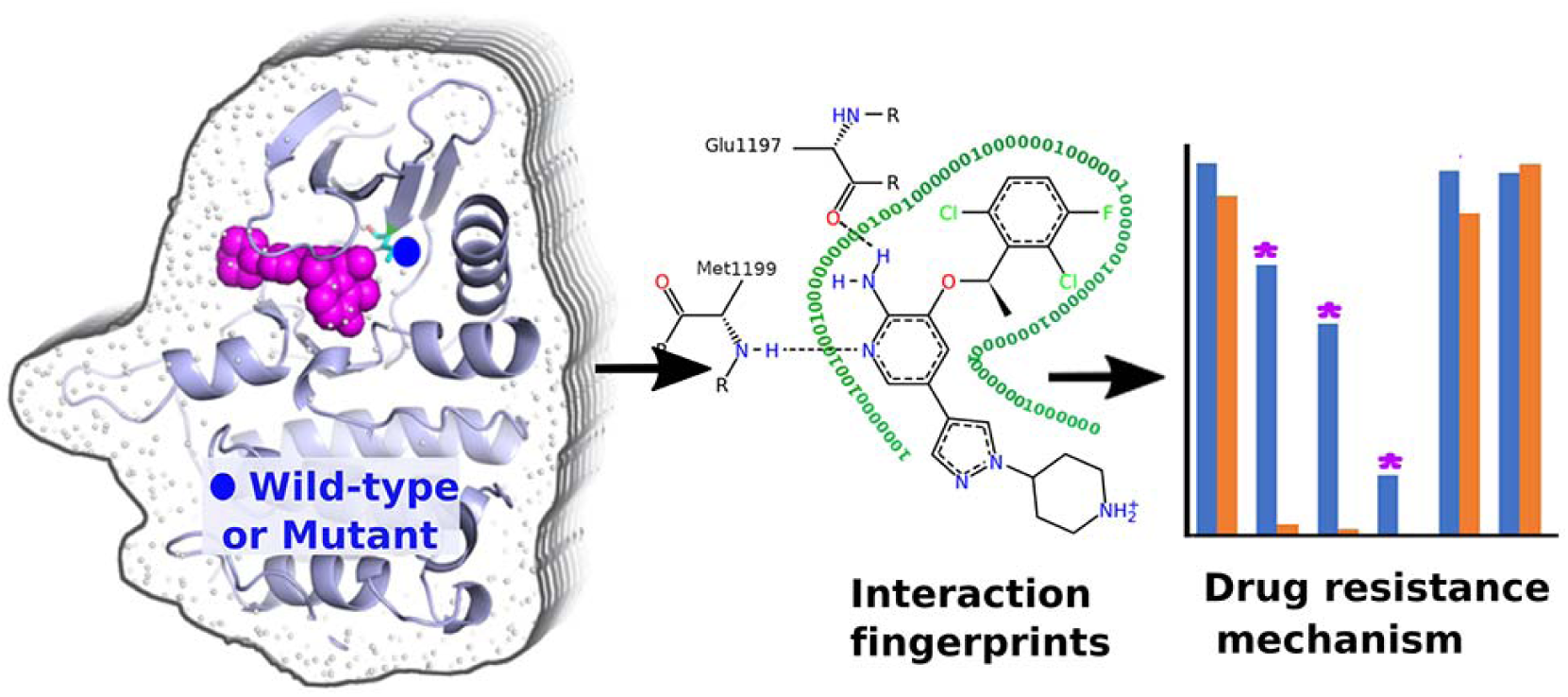

