## Supplemental Figure S1-3 for "Revealing acquired resistance mechanisms of kinase-targeted drugs using an on-the-fly, function-site interaction fingerprint approach"

***Corresponding author**

### **1. Equilibrating the wildtype/mutated drug-bound ALK systems and determining the reaction coordinates**

Before carrying out umbrella sampling for free energy calculations, all ALK systems are equilibrated by running 150ns MD simulations and the corresponding Ca-RMSDs are shown in Figure S1. To determine the common reaction coordinates, we analyzed the respective equilibrated system by calculating the Ca-RMSF of every system (Figure S2). Based on the inflexibility of every residue at the binding site, we chose the rigid residues to build the reaction coordinates (Figure S3). Thus, we chose the distance between the center of mass of the Ca atom of these residues (including residue 1169-1172, 1196-1199, and 1267-1268; green) and the center of mass of ligand core structure (red) as the reaction coordinates to indicate the process of drug binding.

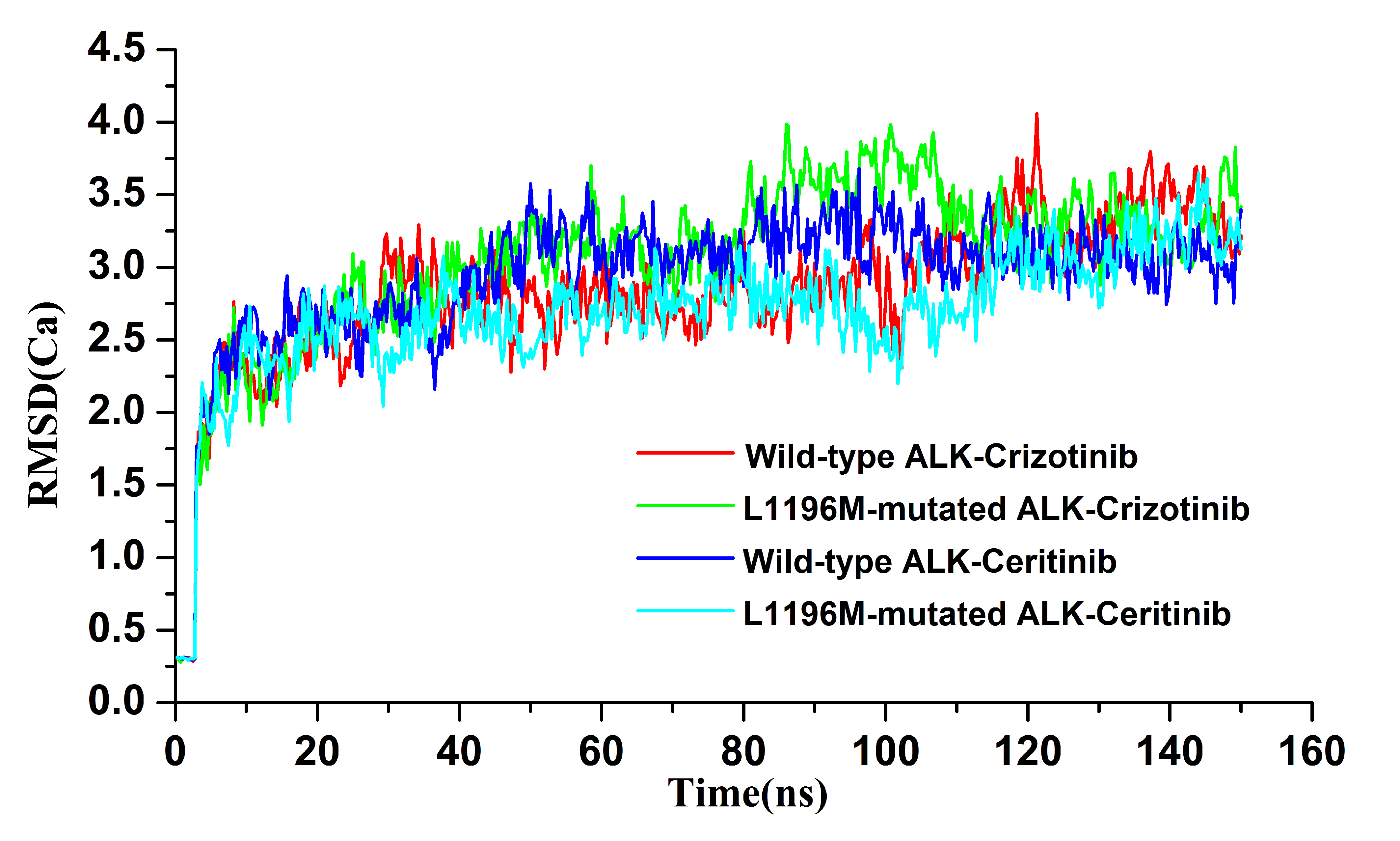

**Figure S1.** Ca-RMSDs of all ALK-drug systems.

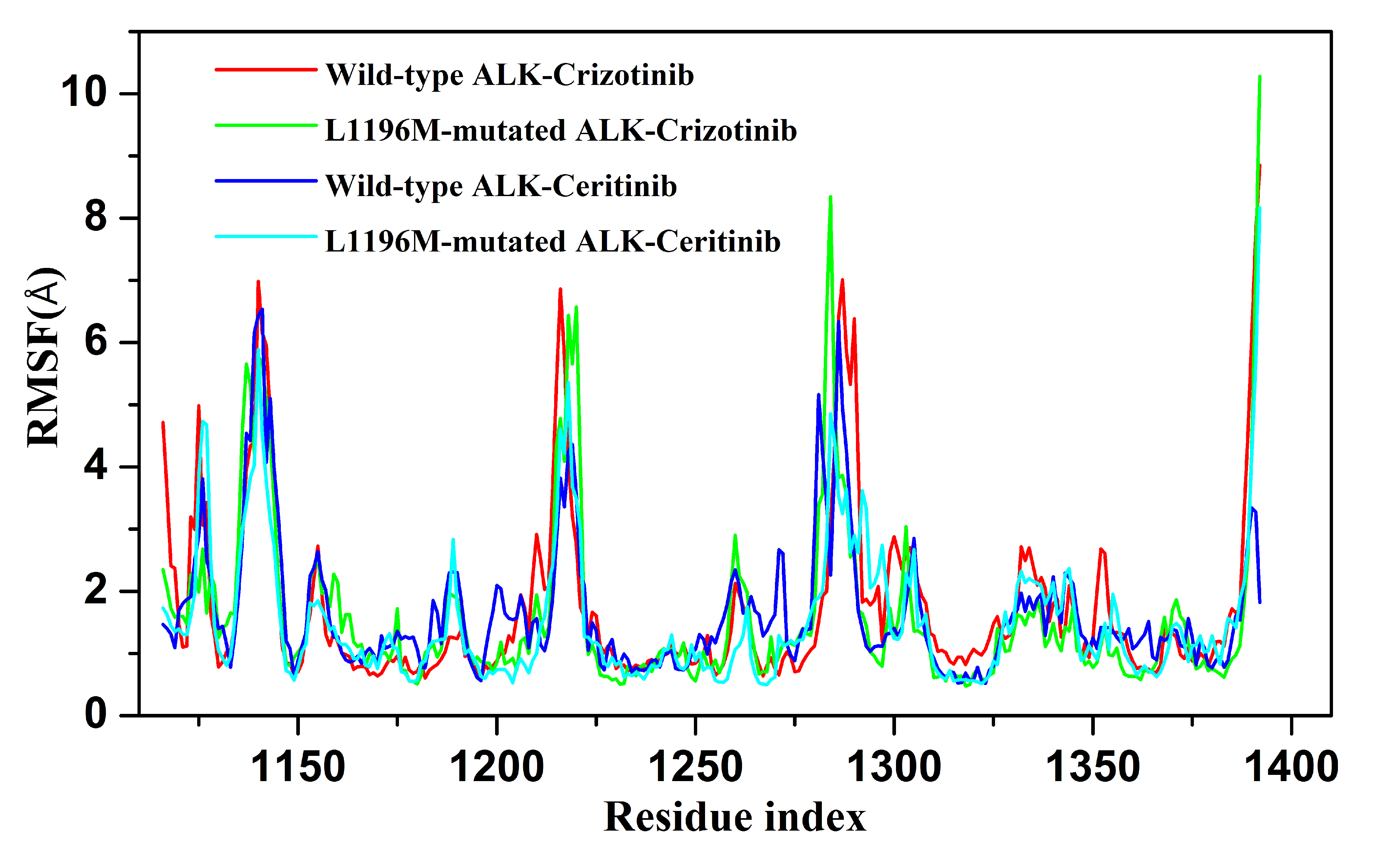

**Figure S2.** Ca-RMSFs of all ALK-drug systems.

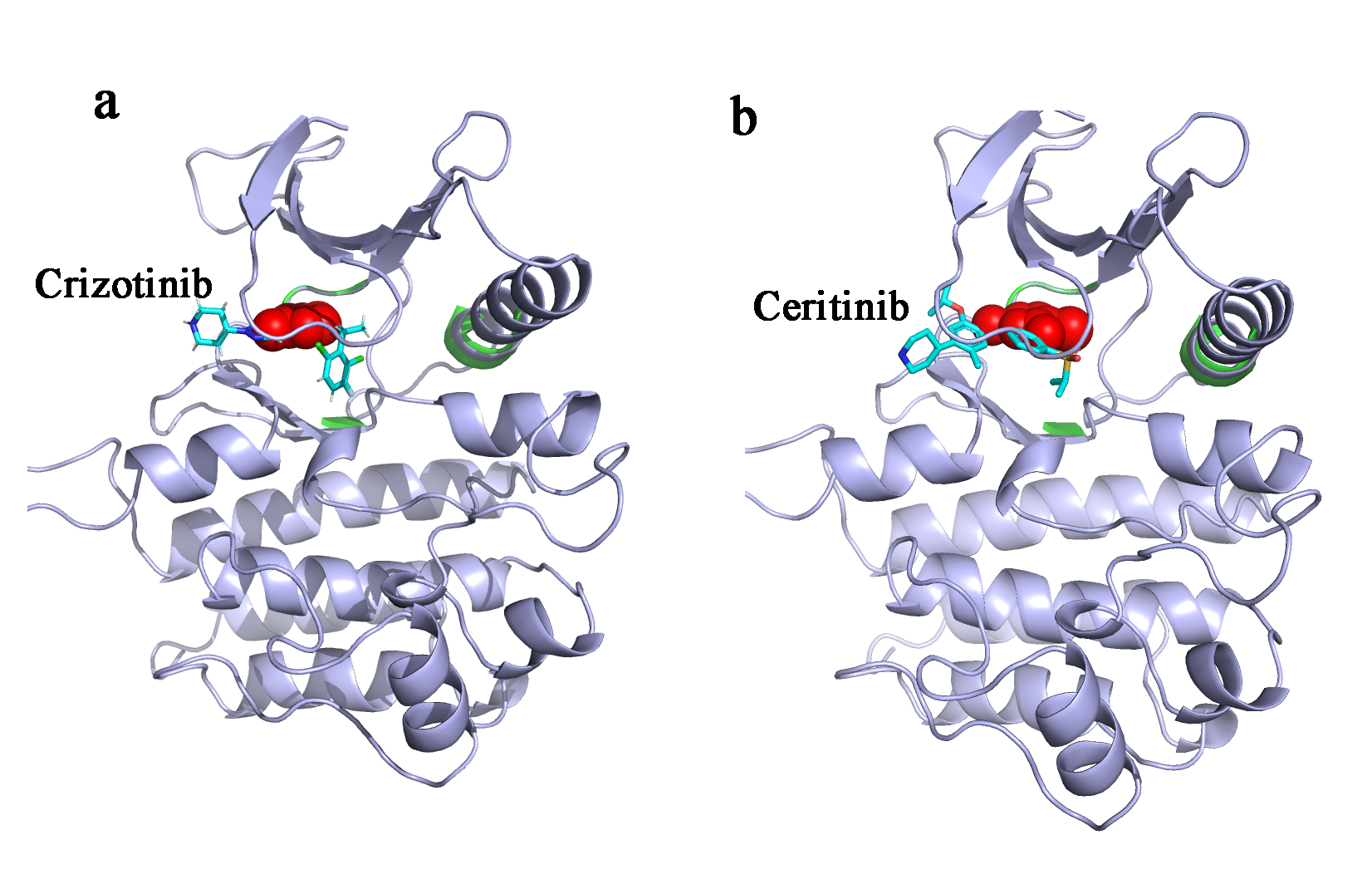

**Figure S3.** Definition of the reaction coordinates for different drug-ALK complexes.

### **2. A brief overview for the WHAM extension**

In umbrella sampling (US) with a biased system Hamiltonian $Hi$,

$Hi=H0+Vi (1).$

$H0$ is the system Hamiltonian and $Vi$ is the bias potential added. The added bias potential will improve the sampling frequency at the fields of high energy. Ideally the $Vi$ should be the negative of potential of mean force $Fi$. $Fi$ is related to Boltzmann distribution, i.e.,

$Fi=-kTln\rho0\left( \lambda\right) (2)$,

where λ is reaction coordinate, and $\rho0$ is the probability distribution. At the beginning the $\rho0$ is not known, however $\rho0$ can be written in terms of non-Boltzmann probabilities $\rho bias$ after a set of biased sampling with bias potential Vi, i.e.,

$\rho0=e^{\left( Vi(\lambda)-Fi \right)/{kBT}}*\rho bias\left( \lambda\right) \left( 3 \right),$

which can be rewritten as:

$e^{{-Fi}/{kBT}}=\int e^{{-Vi(\lambda)}/{kBT}}\cdot\rho0\left( \lambda\right)d\lambda=<e^{{-Vi(\lambda)}/{kBT}}> (4)$.

Where $\rho0$ cannot be calculated directly. The WHAM is an alternative approach designed to get the total probability distribution $\rho0$. The following equations are the WHAM equation. The derivation can be found in the related literatures*^1-3^*.

$$\rho0\left( \lambda\right)=\sum_{i=1}^{N} \frac{ni}{\sum_{j=1}^{N} nje^{{-\left( Vi\left( \lambda\right)-Fi \right)}/{kBT}}}\rho_{i}^{bias}\left( \lambda\right) (5)$$

Where, $ni$ is the number of samples in the $ith$ window. There are non-linear interdependent variables in the equation (4) and (5), which can be solved self-consistently. $Vi$ can be arbitrary function, such as the harmonic restraining formation as we used in the work,

$Vi=ki\cdot\left( \lambda^{'}i-\lambda i \right)2$	(6).

Where $ki=1000 kcal.{mol}^{-1}.Å^{-1}$
